# Z-AAT impairs organelle homeostasis and reduces adaptive response to lipids in alpha-1 antitrypsin deficiency models

**DOI:** 10.64898/2026.08.14.744823

**Authors:** Sara Gil-Martín, Nerea Matamala, Otto Hagen-Doval, Eleonora Bruno, Gema Gómez-Mariano, Carlos Benítez-Buelga, Maria Barrero, Sheila Ramos del Saz, Marta Fernández-Prieto, Selene Martínez, Juliana Manosalva, Diego Megias, Félix Docando, María C. Terrón, Javier Alonso, Antonio Olveira, Miriam Romero, Myriam Calle, Juan Luis Rodríguez-Hermosa, Sabina Janciauskiene, Sara Pérez-Luz, Beatriz Martínez-Delgado

## Abstract

Alpha-1 antitrypsin deficiency (AATD) caused by the Z variant leads to hepatic accumulation of misfolded AAT polymers and liver disease. Although proteotoxic stress is well established, its impact on lipid metabolism, mitochondrial function, and organelle homeostasis remains incompletely understood. The effects of Z-AAT accumulation were investigated in Z-HepG2 cells and 3D patient-derived ZZ hepatic organoids through protein aggregation, lipid storage, mitochondrial structure and function, peroxisomal dynamics, and comprehensive transcriptomic and proteomic analyses. Z-AAT expression led to intracellular polymer accumulation and reduced secretion, together with lipid accumulation, mitochondrial structural abnormalities, increased mitochondrial number but impaired respiratory capacity. Metabolic profiling revealed reduced oxidative phosphorylation and partial reliance on glucose metabolism. Peroxisomes displayed increased mass, consistent with altered lipid handling. Multi-omics analysis demonstrated widespread transcriptional and proteomic reprogramming related to protein synthesis, lipid metabolism, and mitochondrial function. Proteomic analysis confirmed proteotoxic stress-induced mitochondrial dysfunction, impaired lipid handling, and activation of stress response, inflammatory and vesicular trafficking pathways. Importantly, lipid supplementation elicited adaptive mitochondrial transcriptional responses in control cells, whereas Z-HepG2 cells showed a blunted response to lipid challenge. In conclusion, Z-AAT accumulation disrupts hepatic lipid processing and impaired mitochondrial and peroxisomal homeostasis, producing diminished metabolic flexibility likely contributing to AATD-associated liver disease.

## INTRODUCTION

Inherited alpha-1 antitrypsin deficiency (AATD) due to the most common PiZZ (Glu342Lys) genotype is a well-recognized genetic cause of liver disease driven by the accumulation of Z-AAT within hepatocytes (1). Histopathological analysis of PiZZ liver tissue shows characteristic intracellular inclusions that are related to varying degrees of liver damage (2). Metabolic syndrome, a cluster of interrelated metabolic abnormalities including obesity, insulin resistance, hypertension, hypertriglyceridemia, and low HDL cholesterol, is strongly associated with elevated liver aminotransferase levels (ALT, AST, and GGT) as well as an increased risk of hepatic inflammation, fibrosis, and liver injury (3, 4).

The accumulation of misfolded Z-AAT within the endoplasmic reticulum (ER) induces chronic ER stress and activation of the unfolded protein response, which secondarily disrupts hepatic metabolic homeostasis (5–8). Reduced serum triglyceride (TAGs) and lipoprotein levels in individuals with PiZZ, coupled with increased hepatic steatosis, suggest impaired hepatic lipid secretion. Consistent with this, PiZ transgenic mice developed steatosis and exhibited downregulation of lipid export genes (3).

Three-dimensional (3D) organoid cultures that reproduce critical structural and functional characteristics of the human liver provide physiologically relevant models for studying liver diseases (9–11). Patient-derived liver organoids have also recapitulated the key features of AATD-associated liver disease (12, 13). PiZZ patient-derived hepatic organoids, and to a lesser extent PiMZ, but not PiMM organoids display intracellular Z-AAT accumulation, together with increased lipid storage (14). Lipidomic profiling demonstrated a net increase in total lipid content, particularly in TAG, cholesterol esters (core lipid-droplet storage lipids), ceramides (membrane/signaling), and cardiolipins (mitochondrial inner membrane-enriched lipids). These findings provide further support for the altered lipid metabolism in the PiZZ AATD liver.

A consistent finding across species is mitochondrial dysfunction in both PiZZ patient liver biopsies and PiZ transgenic mouse models (15), linking proteotoxic stress to impaired ATP (adenosine triphosphate) production. Consistent with this, studies using AATD patient-derived and genome-edited isogenic iPSC-derived hepatocytes have reported abnormal mitochondrial morphology, reduced membrane potential, and impaired oxidative phosphorylation compared to non-AATD controls (16–18). Functional analyses have further demonstrated reduced basal and maximal respiratory capacity, decreased ATP production, and increased glycolysis, consistent with bioenergetic reprogramming secondary to chronic ER stress and proteotoxic burden (19). More recently, Khodayari et al., using PiZ transgenic mice and Z-AAT-expressing hepatocyte models, suggested that the hepatic accumulation of Z-AAT directly contributes to mitochondrial dysfunction (17).

A better understanding of the molecular pathways driving metabolic liver alterations and mitochondrial injury in AATD is therefore essential to link cellular and biochemical dysfunction with the heterogeneous clinical phenotypes observed in patients with PiZZ. Here, we used human Z-AAT models, including Z-expressing HepG2 cells and ZZ patient-derived hepatic organoids, to better characterize the alterations in lipid metabolism together with morphological and functional changes in metabolism related organelles, mitochondria and peroxisomes. Comprehensive transcriptomic and proteomic analyses allowed us to characterize the interplay among lipid dysregulation, mitochondrial dysfunction, and redox imbalance at the cellular and molecular levels, uncover protein biomarkers of AATD-related liver disease, and identify a lack of metabolic adaptation in Z-AAT-expressing cells in response to lipid challenge.

## RESULTS

### Z-AAT forms intracellular polymers in hepatic cell models

As expected, Z-AAT expression led to pronounced intracellular accumulation and polymerization in both Z-HepG2 cells and patient-derived ZZ-HepORG cells, as shown by immunofluorescence and western blot analyses (Figure 1). In non-Z controls, AAT showed diffuse intracellular distribution with minimal polymer occurrence, whereas Z-HepG2 cells and ZZ-HepORGs exhibited marked perinuclear accumulation of polymeric AAT, consistent with ER retention (Figure 1A). Immunoblot analysis confirmed increased intracellular polymeric Z-AAT and significantly reduced secretion relative to non-Z controls (Figure 1B), validating both models and demonstrating that Z-AAT expression resulted in intracellular polymer accumulation and impaired secretion.

**Figure 1.**
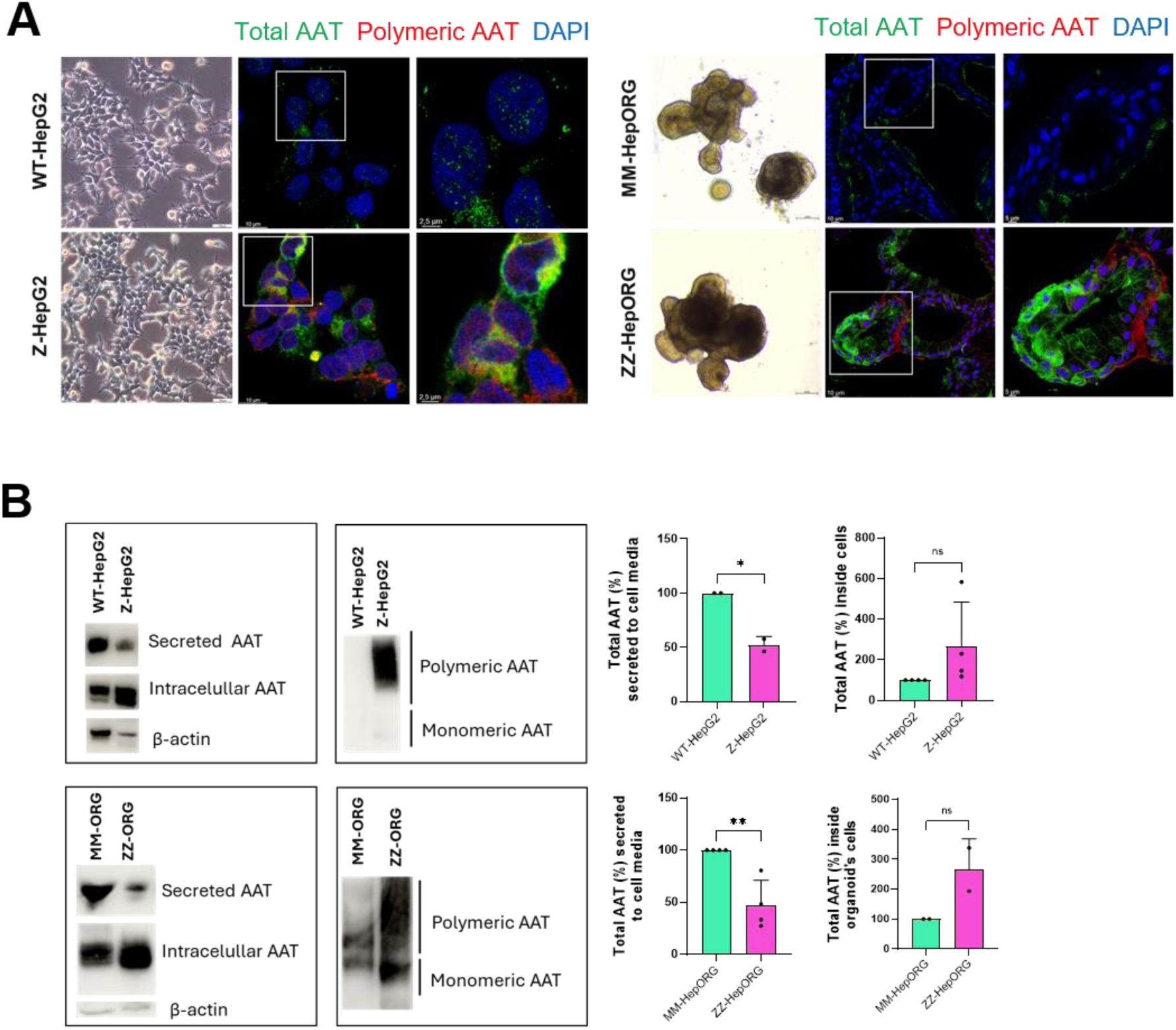
Characterization of Z-AAT expression in cell and organoid models. **A**. Immunofluorescence microscopy of HepG2 cells (left panel) and hepatic organoids (right panel) show an increase in AAT accumulation on both models. Scale bars: 10 µm and 2,5 µm. **B.** Western blot analysis showing a reduction of secreted AAT in Z-HepG2 and ZZ-HepORG (left panels), and the accumulation of polymeric AAT in the insoluble fraction of Z-HepG2 and ZZ-HepORG (right panels).

### Z-AAT expression is associated with increased lipid accumulation

Given that the ER retention of Z-AAT has been linked to metabolic alterations (14, 17), we next assessed intracellular lipid storage. BODIPY staining revealed increased neutral lipid accumulation in Z-HepG2 cells compared to that in WT-HepG2 cells, accompanied by a significant increase in lipid droplet number per cell (Figure 2A). Consistently, Oil Red O staining demonstrated increased neutral lipid deposition in ZZ-HepORGs compared to MM-HepORGs, including focal regions of marked lipid accumulation (Figure 2B). Quantification of perilipin 2 (PLIN2), a lipid droplet-associated protein, confirmed an increase in both the number and size of lipid droplets in Z-HepG2 cells relative to non-Z controls, with a similar but more heterogeneous increase observed in ZZ-HepORGs relative to MM-HepORG (Figure 2C).

**Figure 2.**
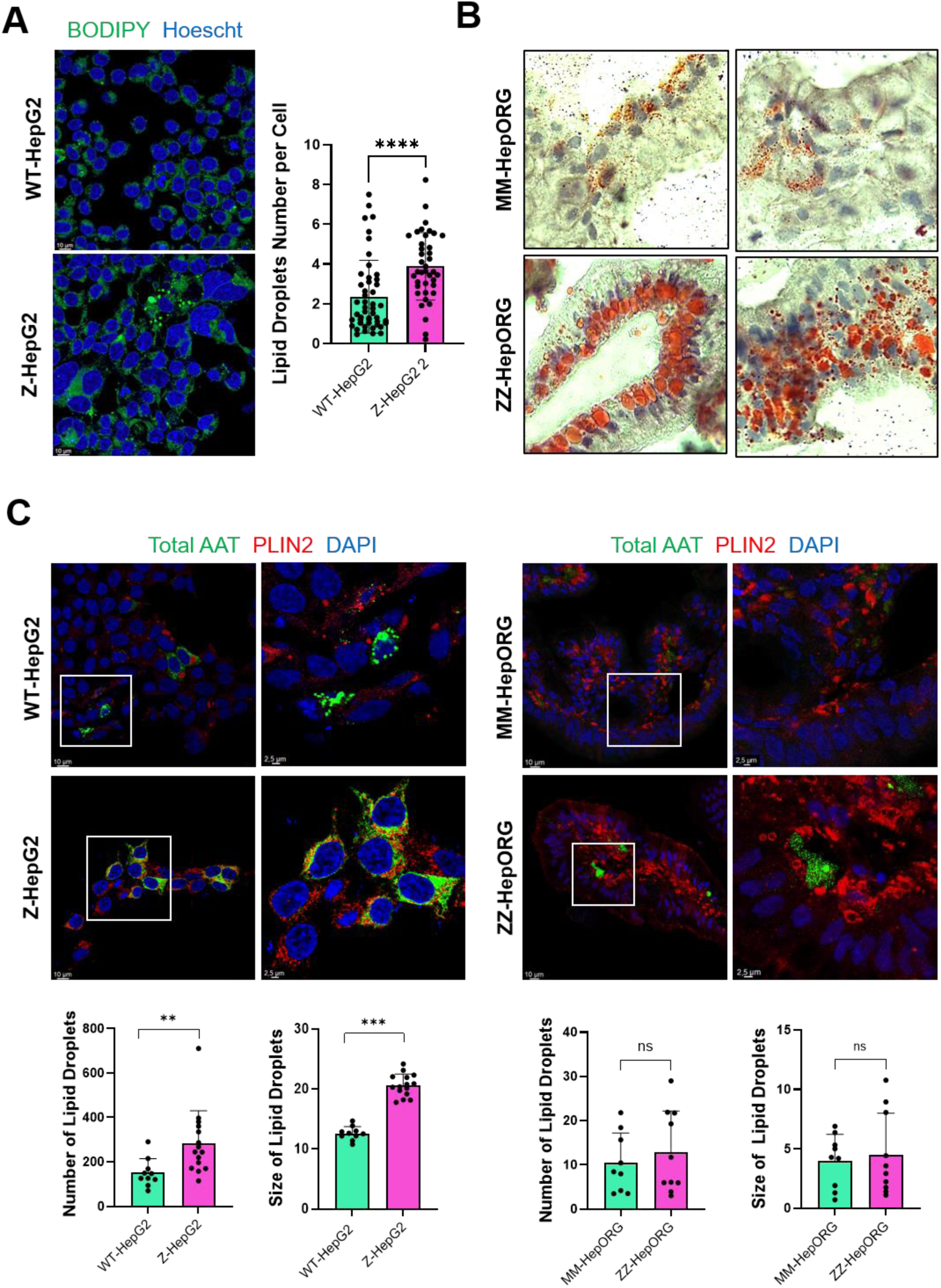
Alterations in lipid storage on Z-HepG2 and ZZ-HepORG. **A.** Immunofluorescence microscopy of BODIPY staining in HepG2 cells displaying increased lipid storage in Z-AAT model. **B.** Immunohistochemistry assay using Oil Red-O showing a bigger lipid buildup on ZZ-HepORG than on MM-HepORG. **C.** Immunofluorescence microscopy of PLIN2 staining (in red) and total AAT (in green) in HepG2 (left panels) and hepatic organoids (right panels) and quantification showing an increase in lipid accumulation more evident in the cell model. Scale bars: 10 µm and 2,5 µm.

### Structural and functional mitochondrial alterations in Z-AAT models

Because of the observed increase in intracellular lipid accumulation, which is often associated with mitochondrial stress and dysfunction, we subsequently investigated the mitochondrial structure and function in Z-HepG2 cells and ZZ-HepORGs. Ultrastructural analysis by transmission electron microscopy revealed marked mitochondrial abnormalities in Z-AAT-expressing cells, including disorganization of cristae architecture and changes in matrix electron density (Figure 3A). Notably, most of mitochondria in Z-HepG2 cells displayed a heterogeneous matrix characterized by areas of reduced electron density, contrasting with the homogeneous and uniformly electron-dense matrix observed in control cells.

**Figure 3.**
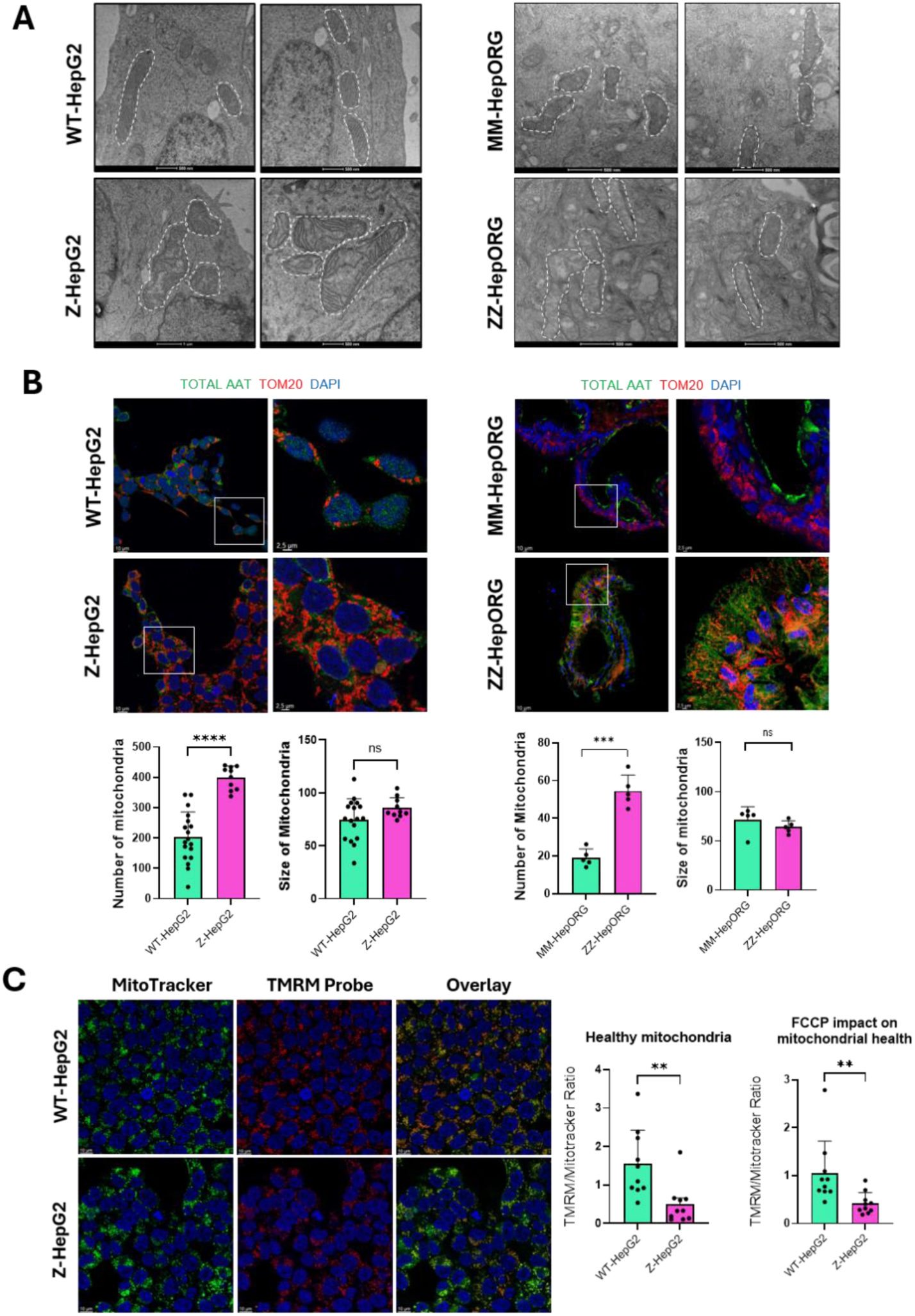
Characterization of mitochondria ultrastructure and function in Z-AAT cells and organoid models. **A.** Representative transmission electron micrographs showing mitochondrial structural alterations in Z-AAT models compared with their respective controls. **B.** Immunofluorescence assay in HepG2 cells (left panels) and hepatic organoids (right panels) using TOM20 as mitochondria marker (in red) and AAT (in green), showing quantification with an increase in mitochondrial number without size alteration. **C.** Live-cell assay of mitochondrial depolarization, using TMRM probe to assess mitochondrial membrane potential and MitoTracker Green to mark mitochondria. Quantification reveals the markedly lower TMRE/MitoTracker ratio in Z-HepG2 cells compared with WT-HepG2 cells (p<0.01) indicating that mitochondria in Z-HepG2 cells are less polarized and therefore have a reduced membrane potential compared with WT-HepG2 cells. This difference was maintained following FCCP treatment, further suggesting impaired mitochondrial functional capacity. Scale bars: 10 µm and 2,5 µm.

To assess mitochondrial content and integrity, we analyzed the outer mitochondrial membrane protein TOM20. Immunofluorescence demonstrated an increased mitochondrial number in both Z-HepG2 cells and ZZ-HepORGs compared to their respective controls, while mitochondrial size remained unchanged (Figure 3B).

We then evaluated mitochondrial function by measuring the membrane potential. Live-cell imaging using TMRM and MitoTracker Green revealed a reduced proportion of polarized (functionally active) mitochondria in Z-HepG2 cells compared to WT controls (Figure 3C). Consistently, the FCCP (uncoupling agent) challenge reduced the ability of Z-HepG2 cells to recover mitochondrial membrane potential, further indicating impaired mitochondrial function in the presence of Z-AAT.

### Mitochondrial metabolic impairment and altered substrate dependency in Z-AAT models

To further investigate the mitochondrial alterations observed in Z-AAT-expressing models, we assessed mitochondrial respiration using the Seahorse XF Cell Mito Stress Test. Z-AAT expression was associated with reduced mitochondrial respiration in both Z-HepG2 cells and ZZ-HepORGs compared with their respective controls (Figure 4A). Z-HepG2 cells showed a trend toward reduced basal respiration and spare respiratory capacity, whereas ZZ-HepORGs exhibited significantly reduced maximal respiration and spare respiratory capacity. Glycolytic function was increased in Z-HepG2 cells compared with control cells, as demonstrated by consistently higher extracellular acidification rates (ECAR), suggesting that Z-HepG2 cells rely more on glycolysis for energy, which implies a shift toward glycolytic metabolism, possibly due to impaired mitochondria (Figure 4A). MM-HepORG engages both mitochondrial respiration and glycolysis, implying its metabolic flexibility. In contrast, ZZ-HepORG is metabolically less active overall or less responsive to energy stress, with both reduced mitochondrial and glycolytic activity.

**Figure 4.**
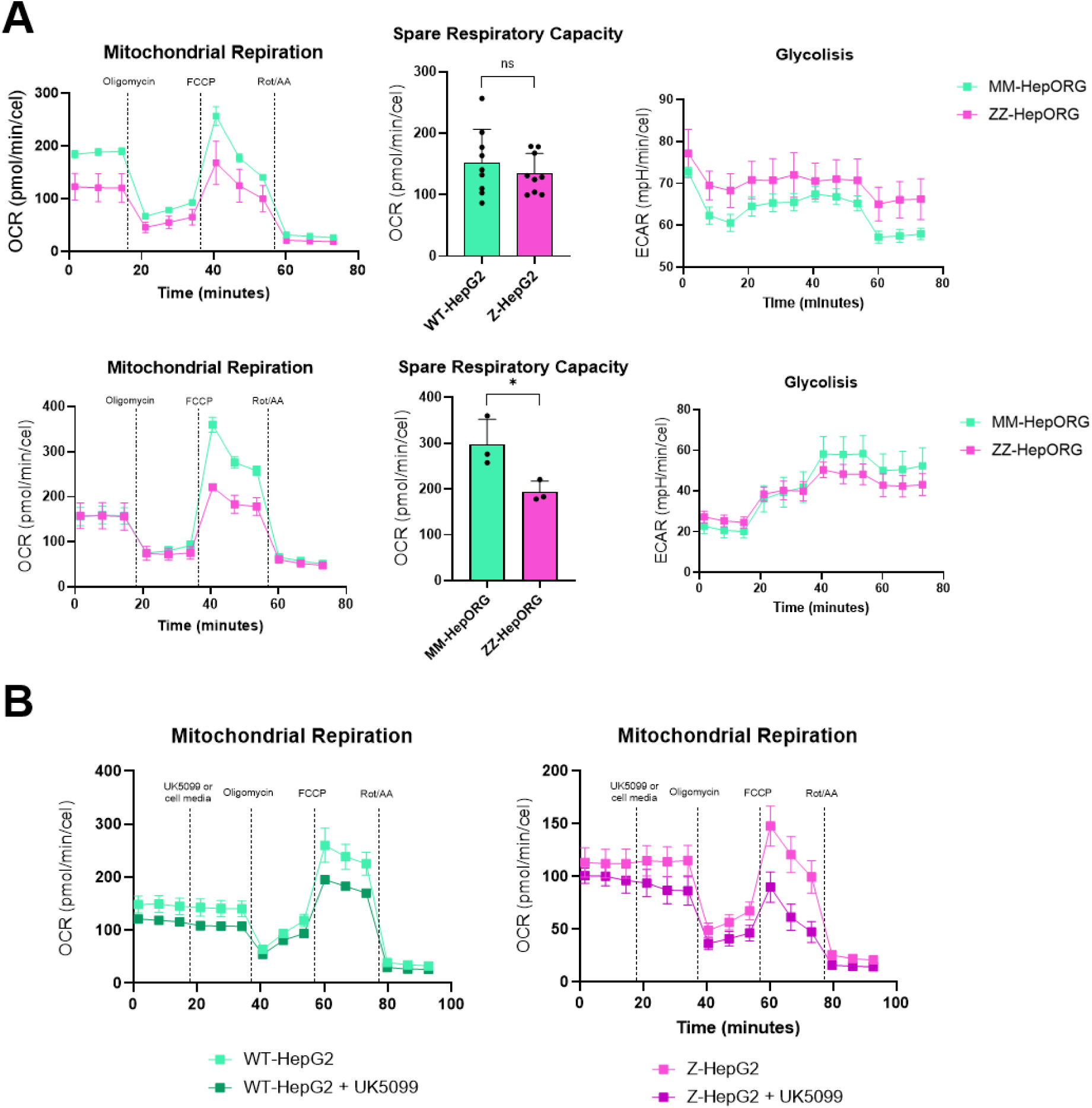
Analysis of the effect of Z-AAT polymers on mitochondrial metabolism. **A**. Mitochondrial respiration represented by the oxygen consumption rate (OCR) (left), differences on spare respiratory capacity in Z-HepG2 and ZZ-HepORG models and their respective controls (center), and glycolysis capacity represented by (ECAR) (right). **B.** Mitochondrial respiration using UK5099 inhibitor of pyruvate import showing that Z-HepG2 cells exhibited a more pronounced reduction in OCR.

To investigate substrate dependency, we inhibited fatty acid oxidation (Etomoxir) and glutaminolysis (BPTES), which did not alter mitochondrial respiration in WT and Z-HepG2 cells (data not shown). In contrast, the inhibition of mitochondrial pyruvate import using UK5099 resulted in a modest decrease in basal respiration, specifically in Z-HepG2 cells (Figure 4B). These findings suggest greater reliance on glucose-derived substrates to sustain mitochondrial respiration in Z-AAT-expressing cells.

### Peroxisomes alteration in Z-AAT models

Peroxisomes are key organelles involved in hepatic lipid homeostasis and contribute to very-long-chain fatty acid β-oxidation and hydrogen peroxide detoxification (20). Given their functional interplay with the mitochondria, we assessed whether Z-AAT accumulation affects peroxisomal morphology and abundance using immunofluorescence for the membrane marker PMP70. In Z-HepG2 cells, the peroxisome number was lower than that in the WT controls, but the organelles appeared enlarged (Figure 5, left panels). In contrast, ZZ-HepORGs showed a significant increase in peroxisome number without changes in size relative to MM HepORG controls (Figure 5, right panels). In HepG2 cells, peroxisome enlargement may reflect increased metabolic demand without the induction of biogenesis. In contrast, patient-derived organoids exhibit increased peroxisome numbers, suggesting the activation of peroxisomal biogenesis pathways and a more coordinated adaptive response. These differences likely reflect variations in metabolic plasticity and organelle crosstalk between simplified cell models and multicellular hepatic systems.

**Figure 5.**
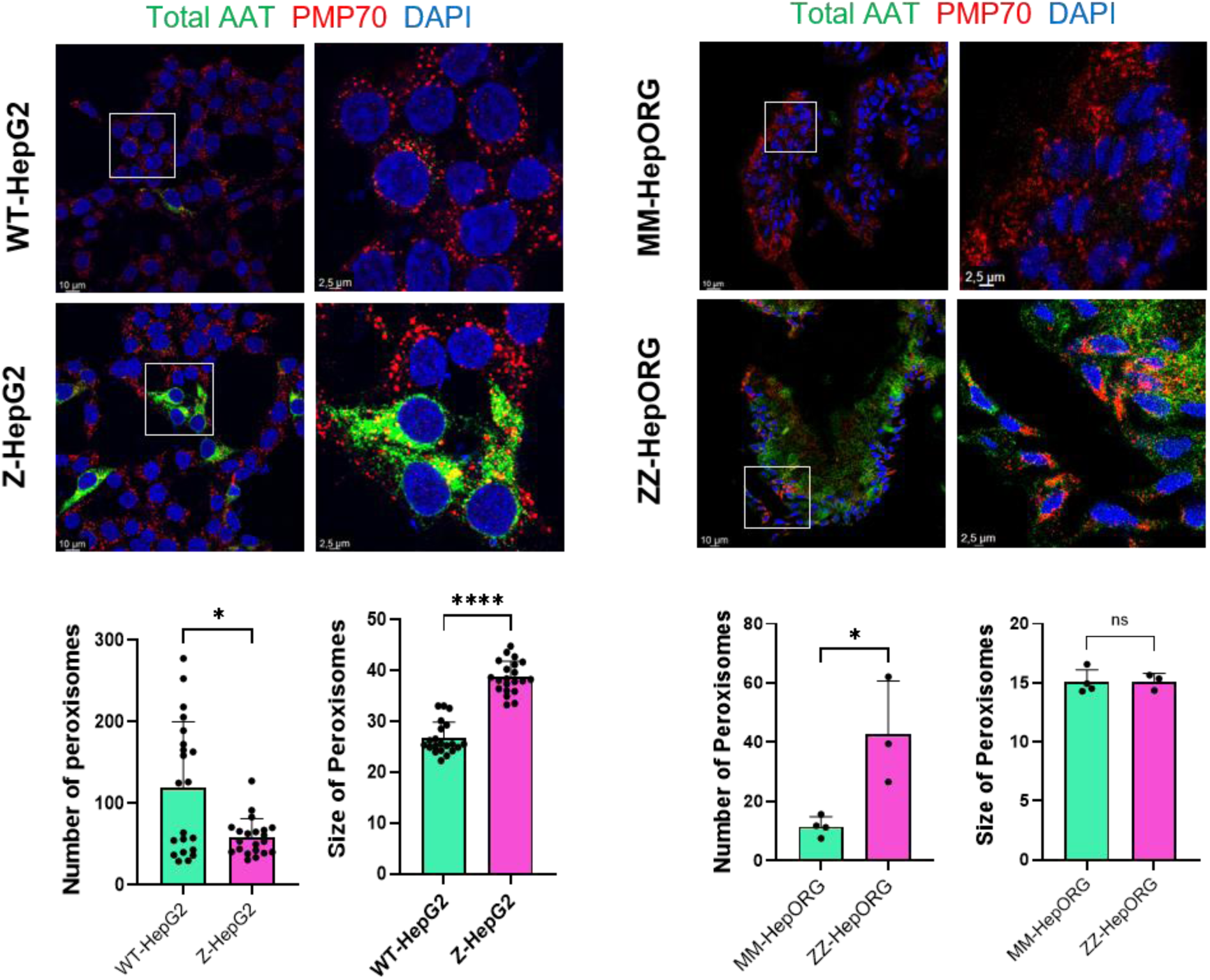
Characterization of peroxisomes in Z-AAT cell and organoid models. Immunofluorescence assay in HepG2 cells (left panels) and hepatic organoids (right panels), using PMP70 (in red) as peroxisome marker and total AAT (in green). Quantification shows a decrease in peroxisome number together with an increase in their size in Z-HepG2 cells, but an increase in peroxisome number in ZZ-HepORG with no change in size. Scale bars: 10 µm and 2,5 µm.

### Z-AAT models exhibit transcriptomic reprogramming consistent with altered lipid metabolism and mitochondrial dysfunction

To further define the extent to which Z-allele expression disrupts cellular homeostasis, we performed transcriptomic profiling of the ZZ-HepORG, MM-HepORG, and Z-HepG2 and WT-HepG2 cells. Principal component analysis of the most variable genes demonstrated clear segregation according to the genotype (Suppl. Figure 1A). Differential expression analysis identified 5208 downregulated and 4110 upregulated genes in ZZ-HepORG compared to those in MM-HepORG (Suppl. Figure 1B). Functional enrichment analysis of differentially expressed genes (DEGs) revealed broad alterations in biological processes associated with protein synthesis, cell cycle regulation, and nucleic acid metabolism (Figure 6A). Gene Ontology analysis further highlighted the significant enrichment of ribosome biogenesis, ribonucleoprotein complex assembly, and translational pathways, indicating global remodeling of cellular protein homeostasis.

**Figure 6.**
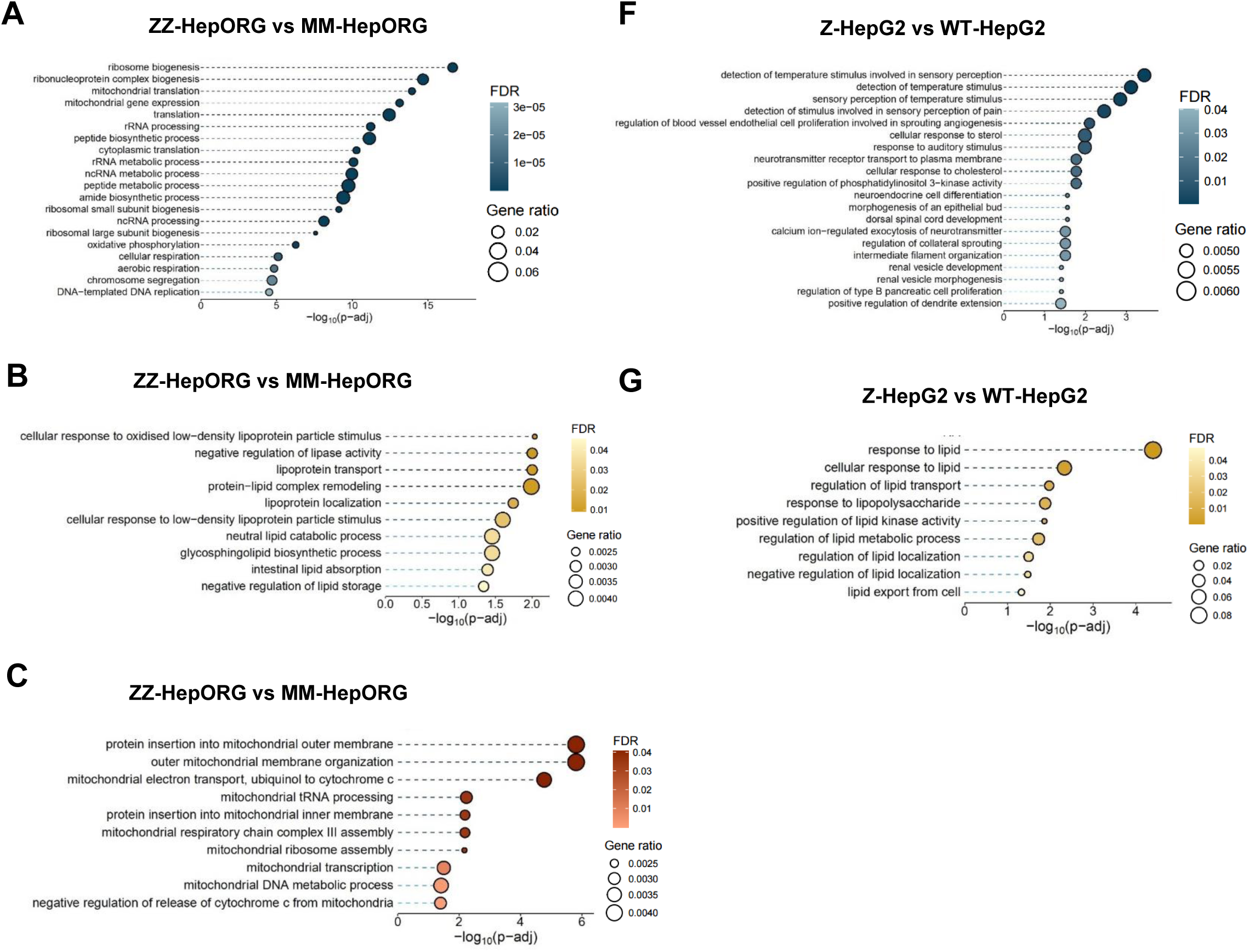
Pathway analysis in the ZZ-HepORG and Z-HepG2 models. **A-C.** Gene ontology biological pathway enrichment analysis (GO:BP) in organoids: top 20 GO:BP terms using all DEG (A), using top 10 biological processes related to lipid homeostasis (B), and top 10 biological processes related to mitochondrial metabolism (C). **D-E.** Gene ontology biological pathway enrichment analysis (GO:BP) in HepG2 cells: top 20 GO:BP terms using all DEG (D) and the top 10 biological processes related to lipid homeostasis (E).

Importantly, analysis of lipid-related pathways demonstrated dysregulation of lipoprotein transport, protein–lipid complex remodeling, neutral lipid catabolism, and glycosphingolipid biosynthesis, as well as responses to LDL and regulation of lipid storage (Figure 6B).

DEGs associated with mitochondrial translation, expression of mitochondrial genes, and mitochondrial ribosome assembly were significantly enriched, indicating broad alterations in mitochondrial protein biosynthesis. Further pathway refinement revealed the dysregulation of mitochondrial transcription, tRNA processing, mitochondrial membrane organization, and impaired assembly of respiratory chain complexes (Figure 6C). Concomitantly, genes associated with lipid metabolism were predominantly downregulated, whereas genes linked to mitochondrial processes were upregulated in ZZ-HepORG compared to MM-HepORG (Suppl. Figure 1C-D).

In Z-HepG2 cells, differential expression analysis identified 316 upregulated and 966 downregulated genes compared to those in WT-HepG2 cells. Enrichment analysis highlighted alterations in pathways related to cellular responses to stimuli along with significant changes in lipid metabolism pathways (Figure 6D-E).

### Proteomic analysis of Z-HepG2 cells reveals metabolic stress and cellular adaptive responses

To complement the transcriptomic findings, we performed proteomic analysis in the Z- and WT-HepG2 models and identified 389 significantly upregulated proteins in Z-HepG2 cells (Figure 7A). Proteomic profiling was not performed in patient-derived organoids because of the limited sample material and the inherent variability associated with patient-derived systems, which can introduce batch effects and reduce quantitative robustness in discovery-based proteomic analyses. Therefore, organoids were primarily used for targeted validation and functional assays.

**Figure 7.**
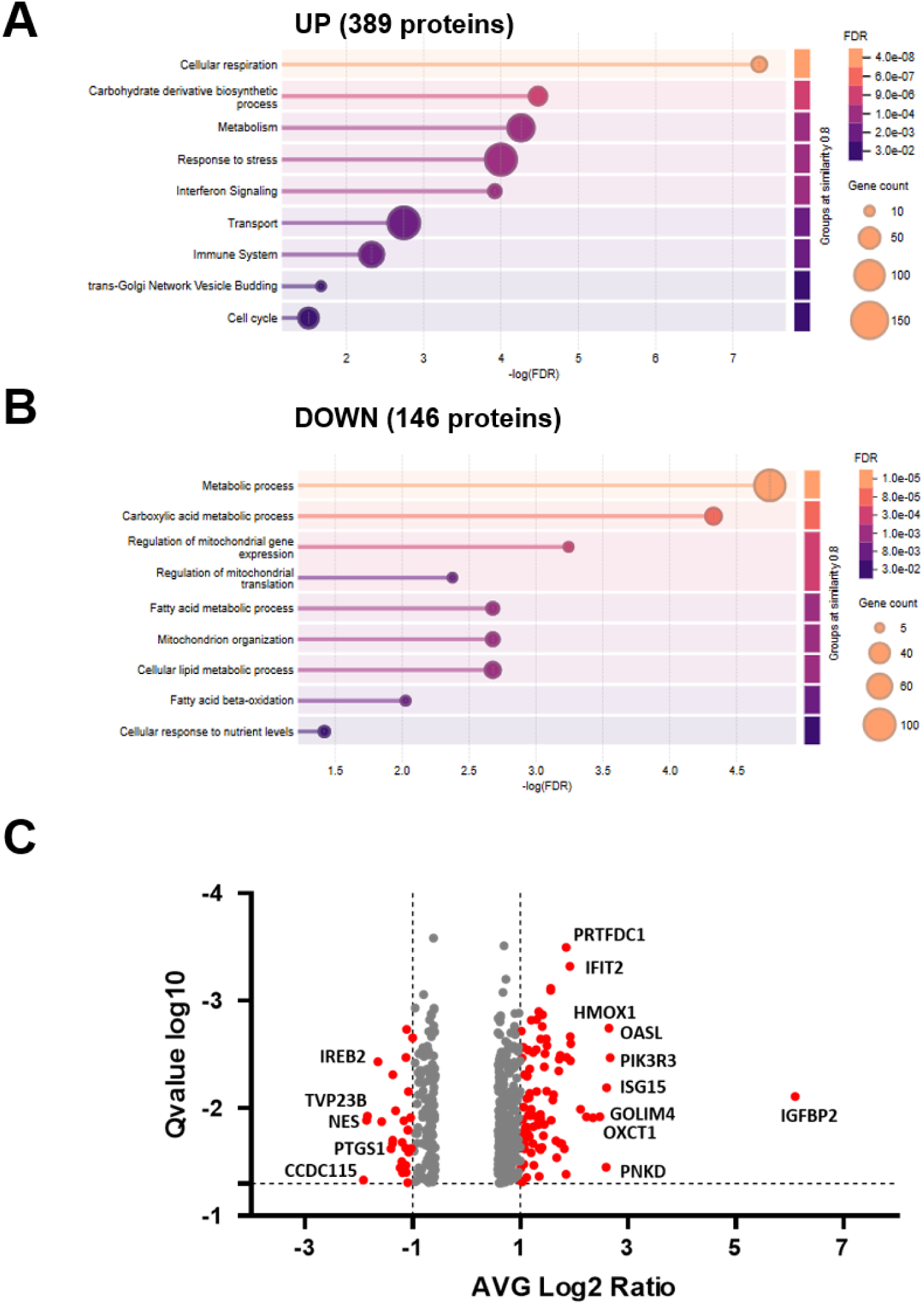
Proteomic changes associated with Z-AAT expression in HepG2 model. Biological processes (GO:BP) enriched in 389 overexpressed proteins (**A**) and 146 downregulated proteins (**B**) in the Z-HepG2 cell model. **C.** Volcano plot with the differentially expressed proteins (q-value<0.05) between Z-HepG2 and WT-HepG2 cells. The x-axis represents the average log₂ fold change in protein abundance (Z-HepG2/WT-HepG2), whereas the y-axis represents the −log₁₀-transformed q-value. Proteins showing a significant increase or decrease greater than 2fold (log₂ fold change ratio >1 or <-1) are highlighted in red. Selected proteins with the largest changes are labelled.

Upregulated proteins were strongly enriched in the mitochondrial pathways, including oxidative phosphorylation, respiratory electron transport, and ATP synthesis. This signature was reflected by the increased abundance of ETC/OXPHOS components (e.g., NDUFA13, NDUFB10, SDHC, UQCRC2, CYC1, COX6C, MT-CO1, MT-ATP6, and MT-ATP8) and ATP synthase subunits (ATP5ME, ATP5MF, ATP5MG, ATP5PB, and ATP5PD). Mitochondrial TOM/TIM protein import machinery (TOMM22, TOMM40, TIMM17B) and metabolite transporters (SLC25A1, SLC25A5) also increased, reflecting the activation of mitochondrial biogenesis and compensatory respiratory adaptation. In addition, Z-HepG2 cells showed a strong upregulation of proteostasis and stress response signatures, including ER stress and autophagy-related proteins (SQSTM1/p62, CALCOCO2, ATF6, HSPD1, UBE2J1), as well as innate immune and interferon signaling activation (DDX58, IFIT1–3, ISG15, STAT1/2, IRF3, and TRIM25). This was accompanied by increased vesicle trafficking and membrane remodeling proteins (RAB5C, RAB11FIP1, SNX1, SEC31A, AP1G2, TMED10) and induction of apoptotic regulators (BAX, CASP8, CASP10) and cell cycle/DNA replication proteins (CDC20, CCNB2, TOP2A, MCM10, RAD51AP1), consistent with a broad stress-adaptive and proliferative response. In contrast, 146 proteins were significantly downregulated and were mainly linked to metabolic dysfunction and organelle impairment (Figure 7B). These included proteins involved in mitochondrial energy production and assembly (COQ8A, SDHAF3, C1QBP, MTO1, TRMU, and MTRES1), suggesting impaired respiratory complex formation, reduced mitochondrial translation, and reduced antioxidant capacity (PRDX2). Levels of enzymes involved in fatty acid β-oxidation and lipid metabolism (ETFDH, ACAD10, ACOX3, DEGS1, and FADS1) were also reduced, consistent with the lipid accumulation observed in the Z-AAT models. Proteins involved in ER–Golgi trafficking and protein processing (PDIA4, DNAJA3, SEC11C, VAPB, SORL1, and LRBA) were decreased, suggesting impaired proteostasis and reduced secretory pathway function. In addition, reduced levels of nutrient- and insulin-signaling regulators (IRS2, SLC38A2, and PRKAG2) could indicate altered metabolic sensing. Finally, decreased expression of autophagy/proteasome and cytoskeletal regulators (PSMD9, WDR45, SEPTIN5, and DYNC2I1) further supports impaired cellular clearance and intracellular trafficking.

Proteins showing the highest changes in Z-HepG2 cells might represent molecular markers of the biological processes taking place in the disease (Figure 7C). They include IGFBP2 as the top upregulated protein, a hepatokine that modulates IGF bioavailability and hepatic lipid/energy metabolism (21). In addition, HMOX1 heme oxygenase-1 is also highly upregulated. HMOX1 catalyzes the first step of heme breakdown into biliverdin, free iron, and carbon monoxide, with antioxidant and anti-inflammatory effects, and therefore it likely stands for a stress-response in the Z-HepG2 cells. Furthermore, PIK3R3 was also one of the top upregulated proteins in Z-HepG2 cells. Overexpression of PIK3R3 in fatty liver models pushes hepatocytes toward fatty-acid oxidation via a PIK3R3-PPARα axis (22). Other upregulated proteins include interferon-stimulated proteins such as OASL, ISG15, IFIT1 and IFIT2, that could represent markers of inflammatory response induced in these cells. On the other hand, the strongest downregulated proteins include CCDC115 (Figure 7C), which acts as an essential factor in Golgi pH-homeostasis, enabling proper protein packaging, and vesicular trafficking as well as glycosylation (23), supporting the altered intracellular transport in Z-HepG2.

### Differential response to lipid supplementation in Z-HepG2 versus WT-HepG2 cells

Finally, we investigated whether lipid supplementation elicited differential transcriptional responses in Z-HepG2 and WT-HepG2 cells (Figure 8A). Gene set enrichment analysis (GSEA) revealed markedly divergent responses between the two cell lines. In WT-HepG2 cells, lipid supplementation induced positive enrichment of mitochondrial and metabolic pathways, including mitochondrial envelope organization, oxidative phosphorylation, respiratory chain complex assembly, and mitochondrial translation. In addition, peroxisomal and proteasomal pathways were positively enriched in response to lipid supplementation, consistent with an adaptive metabolic response aimed at lipid utilization and cellular homeostasis.

**Figure 8.**
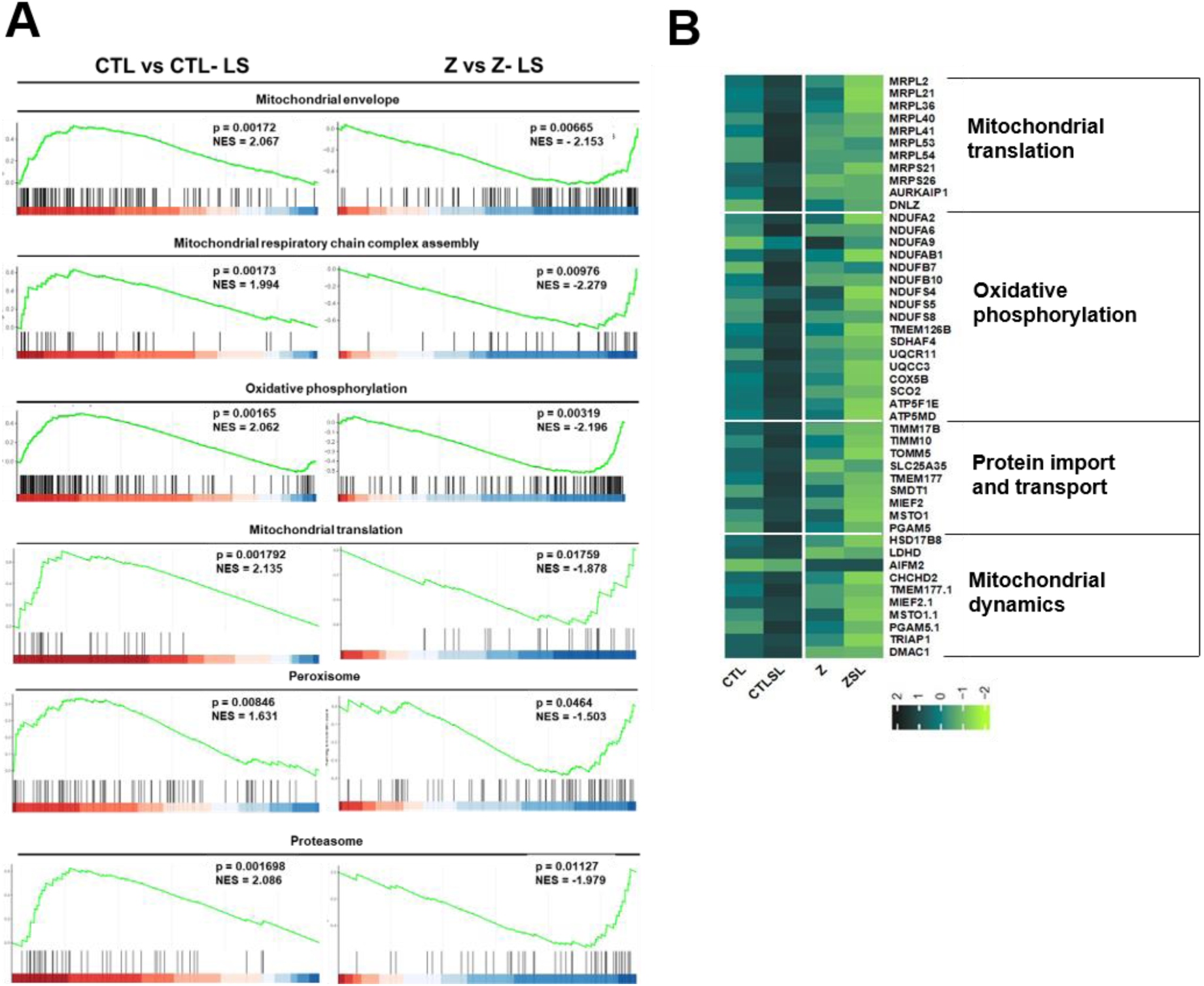
Transcriptomic analysis of the cellular model upon lipid supplement treatment. **A.** GSEA analysis of both wild-type HepG2 and Z-HepG2 cells with and without lipid supplementation (LS). Pre-ranked GSEA analysis shows graphs of positive normalized enrichment scores (NES) in wildtype HepG2 cells (left column) and graphs of negative NES in Z-HepG2 (right column). **B.** Heatmap with the average expression level of mitochondrial-related genes showing opposite behavior in control or Z-expressing cells before and after lipid supplementation. Lower expression of genes is represented in light green and higher expression in dark green.

Z-HepG2 cells displayed the opposite pattern, with significant negative enrichment of the same pathways, indicating downregulation of mitochondrial, lipid metabolism, and protein quality control programs following lipid exposure. This suggests an impaired adaptive response to lipid challenge in Z-AAT-expressing cells, with failure to activate the mitochondrial and metabolic programs required for lipid processing.

To further investigate these differences, we focused on the genes involved in mitochondrial pathways (Figure 8B). In WT-HepG2 cells, lipid supplementation induced upregulation of mitochondrial ribosomal proteins (MRPLs) as well as ribosome-associated factors such as AURKAIP1 and DNLZ, indicating activation of mitochondrial translation. This was accompanied by an increased expression of respiratory chain components and assembly factors, including Complex I subunits (NDUF family genes), UQCR11 (Complex III), COX5B and SCO2 (Complex IV), and ATP synthase subunits (ATP5F1E and ATP5MD). In addition, mitochondrial assembly factors (TMEM126B, UQCC3, and SDHAF4), protein import components (TIMM17B, TIMM10, and TOMM5), and mitochondrial carrier proteins (SLC25A35 and SMDT1) were upregulated, consistent with enhanced mitochondrial biogenesis and oxidative capacity in response to lipid exposure. In contrast, Z-HepG2 cells exhibited reduced expression of mitochondrial ribosomal proteins and respiratory chain components after lipid supplementation. Genes involved in mitochondrial structure and dynamics, including MIEF2 and MSTO1, and stress-response regulators such as PGAM5 and CHCHD2, were also downregulated.

## DISCUSSION

Patients with the classic PiZZ AATD genotype are at increased risk of hepatic diseases (24). Several metabolic abnormalities have been described in PiZZ-associated liver diseases (3, 17, 25). Although mitochondrial impairment and reduced oxidative phosphorylation have been reported, the extent and nature of bioenergetic and organelle-level remodeling in human hepatocytes remain incompletely understood.

In this study, we addressed this gap using human Z-AAT models, including Z-expressing HepG2 cells and patient-derived hepatic organoids (12) to characterize alterations in lipid metabolism, mitochondrial function, and organelle stress responses. Our findings showed that Z-AAT polymer accumulation induces a coordinated multi-organelle stress response characterized by intracellular Z-AAT retention and extensive remodeling of lipid metabolism, mitochondrial function, and peroxisomal homeostasis.

In line with a previous study (14), we demonstrated that Z-AAT accumulation was strongly associated with intracellular lipid accumulation. In Z-HepG2 cells, this is accompanied by coordinated transcriptional and proteomic suppression of lipid metabolism pathways, including fatty acid oxidation and lipid remodeling, suggesting an impaired lipid processing capacity. Importantly, analyses of ZZ-HepORG patient-derived organoids confirmed broader metabolic reprogramming associated with Z-AAT expression. Collectively, these data indicate that the phenotype reflects active metabolic rewiring, rather than a passive consequence of ER retention. Data from clinical studies further supports the role of AATD as a metabolic modifier. In patients with pre-existing metabolic comorbidities, including obesity, alcohol consumption, or metabolic dysfunction-associated steatotic liver disease (MASLD), the presence of Z-AAT aggregates is associated with accelerated progression toward advanced fibrosis and cirrhosis (2), suggesting a synergistic effect between metabolic stress and proteotoxic burden. In parallel, we identified profound mitochondrial alterations at the structural, functional, and transcriptional levels, strengthening previous findings in other animal or cellular AATD models (26). Z-AAT-expressing cells exhibit disrupted mitochondrial ultrastructure, reduced membrane potential, impaired respiratory capacity, and diminished oxidative phosphorylation. These defects are accompanied by reduced expression of proteins involved in mitochondrial maintenance and fatty acid oxidation, together with increased expression of glycolytic cycle system, interferon-response, cell cycle-associated, and mitochondrial structural proteins. Collectively, these data indicate that Z-AAT accumulation induces a coordinated cellular stress response characterized by mitochondrial dysfunction coupled with compensatory metabolic reprogramming. Importantly, integration with our proteomic dataset in Z-HepG2 cells suggested that proteotoxic ER stress may impair mitochondrial translation and lipid oxidation capacity, thereby promoting oxidative stress and metabolic insufficiency. In response, cells appear to engage adaptive programs, including the activation of glycolysis, synthesis of mitochondrial bioenergetic components, autophagy, and innate immune signaling, reflecting an attempt to restore energy homeostasis and maintain viability. Consistent with this model, recent spatial multi-omics analyses of human AATD liver explants have demonstrated that hepatocytes containing Z-AAT aggregates display a profoundly altered proteomic landscape compared with adjacent aggregate-free cells (27). Taken together, these findings support a model in which chronic Z-AAT-driven proteotoxic stress progressively compromises mitochondrial function and limits metabolic flexibility of hepatocytes.

Supporting a direct link between Z-AAT accumulation and mitochondrial dysfunction, RNA interference (RNAi) therapeutics in PiZ mouse models have been shown to reduce the Z-AAT polymer burden, improve mitochondrial homeostasis, and attenuate liver injury (28). Moreover, a previous study suggested that misfolded Z-AAT might be directly associated with mitochondria, thereby contributing to respiratory chain impairment and altered lipid metabolism (17). In line with this, our multi-omics analysis identified dysregulation of mitochondrial translation, respiratory chain assembly, and protein import pathways, indicating that Z-AAT-associated mitochondrial dysfunction occurs across multiple levels of organelle biogenesis and maintenance. Notably, mitochondrial impairment was not constitutive but became particularly evident under metabolic stress. Upon lipid supplementation, control cells activate a coordinated adaptive program involving mitochondrial biogenesis, oxidative phosphorylation, and peroxisomal metabolism, reflecting efficient metabolic adaptation to increased lipid availability. In contrast, Z-AAT-expressing cells failed to engage in this adaptive response and instead displayed suppression of mitochondrial and lipid metabolism transcriptional programs, revealing a strong defect in the adaptive response in response to lipids.

The inability of Z-HepG2 cells to induce mitochondrial translation and respiratory programs in response to a lipid challenge may represent a key mechanism linking Z-AAT proteotoxicity to hepatocellular lipid accumulation. Instead of increasing the oxidative capacity to buffer excess lipid load, Z-AAT-expressing cells remain in a state of mitochondrial insufficiency, which may promote incomplete fatty acid oxidation, reactive oxygen species production, and progressive lipid droplet accumulation. Together, these findings identify impaired transcriptional metabolic adaptability as a previously under-recognized feature of AATD hepatocytes that may contribute to disease progression under conditions of increased lipid availability.

Our findings also extend to peroxisomal remodeling, where we observed increased peroxisomal mass in the Z-AAT models, likely to reflect a compensatory response to lipid overload and oxidative stress. Given the close functional interplay between peroxisomes and mitochondria in fatty acid oxidation and redox homeostasis, these alterations further support the concept of integrated organelle stress in AATD hepatocytes.

In parallel, proteomic profiling of Z-HepG2 cells uncovered a set of previously unreported proteins associated with Z-AAT overexpression, such as IGFBP2, PIK3R3, HMOX1 indicative of the coordinated remodeling of cellular stress-adaptation networks. Other proteins are linked to mitochondrial function, proteostasis, vesicular trafficking, and inflammatory signaling, suggesting that Z-AAT accumulation induces systems-level reorganization of hepatocellular homeostasis beyond lipid metabolism alone. Therefore, these proteins might represent early adaptive regulators and potential mechanistic mediators connecting proteotoxic stress to metabolic dysfunction in AATD liver disease, that remain to be further explored in other systems.

Altogether, transcriptomic and proteomic analyses support a model in which Z-AAT accumulation drives a broad stress response involving proteostasis, autophagy-related pathways and altered vesicular trafficking. Activation of inflammatory signaling further suggests engagement of innate immune and cellular stress programs beyond metabolic dysregulation alone. Additional patient-derived data further support this concept, indicating that chronic Z-AAT retention drives a multi-organelle cascade that contributes to liver injury. A characteristic histological hallmark of PiZZ AATD is the presence of PAS-D-positive hepatocytic globules, which represent Z-AAT polymers retained within the rough ER. Persistent ER retention overwhelms cellular quality control systems, including ER-associated degradation and autophagy, resulting in sustained activation of the unfolded protein response, inflammation, hepatocyte injury, and progressive fibrosis (29).

Overall, our findings support a model in which Z-AAT accumulation drives a coordinated breakdown of hepatocellular homeostasis across the ER, lipid, mitochondrial, and peroxisomal networks. Instead of affecting isolated pathways, Z-AAT proteotoxicity induces an integrated organelle stress response that compromises metabolic flexibility and impairs the ability of hepatocytes to adapt to lipid challenge. This failure of adaptive capacity results in persistent organelle alterations, mitochondrial insufficiency, dysregulated lipid handling, and sustained cellular stress, thereby promoting progressive liver injury in AATD patients.

## METHODS

### Generation of a stable Z-allele expressing cell line

A stable HepG2 cell line expressing the Z-allele (hereafter referred as Z-HepG2) was generated by cloning the mutated *SERPINA1* cDNA into the pRRL.sin.cPPT.CMV.Wpre lentiviral plasmid (30). Briefly, the Z allele was extracted from the plasmid pCMV6 (Origene) by digestion with *Nde*I and *Sma*I restriction enzymes. On the other hand, pRRL.sin.cPPT.CMV.Wpre (hereafter referred to as pLV vector) was initially digested with *Kpn*I, followed by purification and treatment with T4 polymerase enzyme to remove sticky ends, and then digested with *Nde*I enzyme. Both fragments were ligated using T4 ligase to obtain the pLV-Z vector.

Lentiviral particles containing a lentiviral vector with the Z allele were produced by transient transfection of human embryonic kidney 293T cells with Lipofectamine (Invitrogen Life Technologies) and Plus Reagent (Invitrogen) following the manufacturer’s recommendations. Each lentiviral construct (5 μg) was co-transfected with 5 μg of the packaging plasmid, pCMVdR8.74 Addgene plasmid 22036, providing all vector proteins driven by the hCMV promoter but the envelope protein, and 2 μg of the plasmid pMD.2G Addgene plasmid 12259, encoding the heterologous vesicular stomatitis virus envelope, in a p100 plate previously seeded with 2 × 106 cells. The viral supernatant was collected 48 hours after transfection. The titer of the obtained supernatant was determined by flow cytometry using the GFP protein present in the vector as a reporter, as previously described (31). Virus-containing supernatant was used to transduce HepG2 cells, and 24 h post-infection, the cells were seeded by limiting dilution to obtain clonal cultures. The cultures were carefully examined for the appearance of colonies. After the cells reached confluence, they were trypsinized and expanded for further characterization.

### Generation and culture of patient-derived hepatic organoids

Hepatic organoids were established from liver biopsies obtained from patients and control individuals at the Hospital Universitario 12 de Octubre (Madrid, Spain), and undifferentiated adult progenitor cells from established organoid cultures were subsequently deposited at the BioNER biobank at ISCIII for use in further research. In this study, ZZ organoids were generated from two patients with AATD carrying the PiZZ genotype who underwent liver transplantation due to end-stage liver disease (Table 1). Control MM organoids were established from histologically normal liver tissue obtained from two individuals undergoing surgical resection for hepatocellular carcinoma, specifically selected based on the absence or low hepatic steatosis to avoid potential confounding effects of lipid accumulation.

**Table 1.** Clinical characteristics of the patients used to establish organoid lines.

| Genotype | Sex | Age<br>(Years) | Surgery | Serum AAT<br>(g/L) | ALT<br>(U/l)<br>(5-45)* | AST<br>(U/l)<br>(5-33)* | GGT<br>(U/l)<br>(8-61)* | Platelets<br>(x1000/μl)<br>(140-450)* | Glucose<br>(mg/dl)<br>(70-110)* | Albumin<br>(g/dl)<br>(3,5-5)* | Bilirubin<br>(mg/dl)<br>(0,2-1)* |
| --- | --- | --- | --- | --- | --- | --- | --- | --- | --- | --- | --- |
| MM | F | 78 | Hepatocellular carcinoma | ND | 31 | 31 | 49 | 246 | 113 | 4,5 | 0,3 |
| MM | F | 67 | Hepatocellular carcinoma | ND | 43 | 32 | 16 | 265 | 75 | 3,9 | 0,8 |
| ZZ | M | 13 | Neonatal Cholestasis.<br>Liver transplant | 0,31 | 60 | 66 | 25 | 67 | 83 | 3,7 | 1,36 |
| ZZ | M | 1 | Hepatic failure.<br>Liver transplant | 0,41 | 199 | 360 | 551 | 92 | 71 | 3 | 3,67 |

\* Range of normal values; ND: not determined.

Liver tissue biopsies were collected and processed to derive hepatic MM and ZZ organoids as previously described (12). Organoids were expanded in basement membrane extract type 2 (BME-2) using expansion medium (EM). To induce hepatocyte differentiation, organoids were switched to a differentiation medium (DM) and maintained under these conditions for 15 days (32). Differentiated hepato-organoids from MM and ZZ-AATD patients showed no morphological differences and confirmed to express typical mature hepatocyte markers (12). When needed, organoids were collected and dissociated either manually or using TrypLE Express 1X (Gibco) according to the assay they were needed for.

### RNA extraction

Total RNA from cell lines was purified with the RNeasy Mini Kit (Qiagen) following the manufacturer’s recommendations whereas RNA from organoids was extracted using the TRIzol Reagent (Life Technologies) followed by DNase I digestion to avoid contamination with genomic DNA. RNA concentration and purity were measured using NanoDrop 2000 (Thermo Fisher Scientific), and integrity was assessed using the RNA Nano 6000 Assay Kit of the Agilent Bioanalyzer 2100 system (Agilent Technologies).

### Transcriptomic analysis (RNASeq) of the stable Z-HepG2 cell line

For RNASeq library preparation of WT and Z-HepG2 cells, the Hieff NGS Ultima Dual-mode mRNA Library Prep Kit for Illumina was used following the manufacturer’s recommendations. Each cell line was analyzed in triplicates. Poly(A)-containing mRNA molecules were purified using poly(T) oligo-attached magnetic beads. Library quality control was performed using QSep-400 (BiOptic) and Qubit 3.0 (Thermo Fisher Scientific). The qualified library was sequenced on the Illumina Novaseq X platform (Illumina) in paired-end 150 bp (PE150) mode. Library construction and sequencing were performed at Biomarker Technologies (BMKGENE) GmbH.

RNASeq data in FASTQ format were mapped against the reference human genome GRCh38_release95.genome.fa. using HISAT2 (33) and StringTie (34) and the alignment quality was assessed using RseQC v3.0.1 (Wang et al., 2012). Quantification of gene expression levels was estimated by fragments per kilobase of transcript per million fragments mapped (FPKM). Bioinformatics analysis was conducted using R and the BMK Cloud platform (www.biocloud.net). Differentially expressed genes (DEGs) between the Z-AAT models and their respective controls were identified using edgeR (35). The criteria for differentially expressed genes were set as Fold Change (FC) ≥ 1.5 and Pvalue < 0.05. Functional enrichment analysis was performed using Gene Ontology (GO) (adjusted p-value < 0.05) to identify biological processes affected by the ZZ genotype.

### Transcriptomic analysis (RNASeq) of organoids

For RNA-seq, organoids from controls and organoids derived from two ZZ patients were analyzed in triplicate. Library preparation was performed using the TruSeq Stranded mRNA Kit (Illumina) according to the manufactureŕs recommendations. Sequencing was performed at the Genomics Unit (ISCIII) on a NovaSeq 6000 sequencer using 100 base read lengths in paired-end mode.

RNA sequencing data in FASTQ format were analyzed using the nf-core/rnaseq pipeline v3.10.1 (https://github.com/nf-core/rnaseq/releases/tag/3.10.1), an RNA-seq workflow developed by the nf-core community (https://nf-co.re/) and implemented in Nextflow (https://www.nextflow.io/).

Raw reads were initially evaluated for quality using FastQC v0.11.9 (http://www.bioinformatics.babraham.ac.uk/projects/fastqc/). Adapter trimming and removal of low-quality bases at the 3′ ends were performed with TrimGalore v0.6.7 (https://github.com/FelixKrueger/TrimGalore). High-quality reads were then aligned to the ENSEMBL human reference genome GRCh38 using STAR v2.7.9a (36), and the alignment quality was assessed using RseQC v3.0. (37). Transcript quantification was carried out using the featureCounts function from the Subread package v2.0.1 (38).

Downstream analyses were performed using R v4.3.3 (*R: The R Project for Statistical Computing*, s. f.). Lowly expressed genes were filtered out when they had fewer than 15 read counts across all samples. Differential expression analysis was carried out using DESeq2 v1.42.1 (39) with the design formula ∼ Genotype, a factor with two different levels (i.e., MM, ZZ). Genes were considered differentially expressed (DEG) using a 5% FDR and an absolute log2 Fold Change > 0 as thresholds. Gene Ontology (GO) overrepresentation analysis of biological process terms was performed using clusterProfiler v4.10.1 (40) and org.Hs.eg.db v3.18.0.

### Proteomic analysis

Proteins extracted in RIPA buffer from each cell line in triplicate samples, were reduced with 100 mM TCEP (45 min, 37°C) and alkylated with 0.4 M chloroacetamide (30 min, RT, dark). Protein cleanup and digestion were performed using SeraMag magnetic beads (1:1 hydrophilic/hydrophobic) with ACN-assisted binding, followed by washing and overnight digestion with porcine trypsin in 50 mM ammonium bicarbonate (pH 8.0). The peptides were dried under vacuum and stored at −80°C until analysis.

LC–MS/MS was performed on an Orbitrap Astral mass spectrometer coupled to a Vanquish Neo UHPLC (Thermo Fisher Scientific). Peptides were trapped on a PepMap Trap cartridge and separated on an Easy-Spray PepMap RSLC C18 column (75 µm × 15 cm) at 50°C using a 15 min gradient at 300 nL/min. Data were acquired in the DIA mode over m/z 380–980 using 299 variable windows (2 m/z isolation, NCE 25). Full MS scans were acquired at 240,000 resolution.

Raw data were analyzed with Spectronaut (v20.2.250922) using directDIA and UniProt human database (UP000005640_9606, June 2025). Trypsin/P and Lys/P were specified with up to two missed cleavages; carbamidomethylation of cysteines was set as a fixed modification, and methionine oxidation and N-terminal acetylation were set as variable modifications. Identification was filtered at 1% FDR at peptide and protein levels, with IDPicker used for protein inference, and no imputation applied. Differential expression analysis was performed defining differentially expressed genes at 5% FDR and |log2 fold change| > 0.5. Gene ontology overrepresentation analysis of biological processes was conducted using STRING (41).

### AAT detection and quantification by Western Blot

To detect monomeric and polymeric AAT, proteins from the cellular pellets and media were extracted and separated by SDS-PAGE. Protein concentration was quantified using the PIERCE BCA Protein Assay Kit (Thermo Fisher Scientific) following the manufacturer’s recommendations. Native and polymeric forms of AAT were detected, as previously described (12) using either denaturing or native polyacrylamide gels.

The primary antibodies used were rabbit anti-AAT (DAKO, #A0012) at 1/1000 for detection of total AAT, mouse ATZ11 (42)) at 1/500 for detection of polymeric AAT, and mouse anti-β-actin (A1978, Sigma Aldrich) at 1/1000 dilution; the secondary antibodies used were sheep anti-mouse IgG-HRP (NA931V, Heatlhcare) and donkey anti-rabbit (NA934V, Heatlhcare), both at 1:5000 dilution. Quantification of western blot bands of AAT protein was carried out using ImageJ software v1.48, using β-actin (housekeeping protein) to normalize the band’s intensity.

### Oil Red-O Staining

To detect lipid accumulation and lipid droplets, organoid slices embedded in gelatin were fixed with 4% PFA and stained with Oil Red O (Sigma-Aldrich) following manufacturer’s recommendations.

Afterwards, the organoid slices were washed and stained with Mayer’s hematoxylin. Eukkit (Chem-Lab NV) or glycerol 60% was used as a mounting medium. Finally, the stained organoid slices were scanned using a NanoZoomer-SQ Digital slide scanner (Hamamatsu), and the intracellular lipid content was quantified using ImageJ software and CellProfiler.

### Immunofluorescence assays

Cells were seeded at a specific density, allowing them to reach 80% confluence the day of the assay. They were grown either on 8 well plates (Ibidi) to carry out *in vivo* imaging or on top of 12 mm coverslips placed on M6 well plates. Cells grown on M6 plates were fixed with 4% PFA for 15 min, and after washing three times with PBS 1X, the blocking solution (PBS 1X, Triton X100 0,25% and NHS 5%) was added to the wells. The same protocol was followed for staining gelatin slices of the differentiated hepatic organoids.

A combination of different markers was used for each assay: rabbit anti-AAT (DAKO, #A0012) at 1/500 or mouse B9 (SantaCruz, #sc59438) at 1/100 was used for detection of total AAT, mouse D11 (ATZ11) at 1/100 was used for detection of polymeric AAT, mouse Tom20 (SantaCruz, #Sc-17764) at 1/250 was used for detection of mitochondria, rabbit Plin2 (Proteintech, #15294-1-AP) at 1/500 was used for the detection of lipid droplets, and rabbit PMP70 (Abcam, #ab3421) at 1/500 was used for peroxisome detection. A combination of Alexa Fluor 488 (Invitrogen) and Alexa Fluor 594 (Invitrogen) at 1:500 dilution was used as secondary antibodies.

To carry out immunoassays on live cells, they were incubated for 15–30 min with a specific marker diluted in DMEM without phenol red (Corning) for each assay. Afterwards, they were incubated with Hoechst (Invitrogen, H3570) at a 1/300 dilution and immediately observed by immunofluorescence microscopy at 37°C. To assess the mitochondrial membrane potential, HepG2 cells were incubated with MitoTracker Green (M7514, Invitrogen,) at a final concentration of 1 µM. To assess mitochondrial capacity for recovery, HepG2 cells were incubated with TMRM Probe (Invitrogen, I34361) at a final concentration of 0,5 µM, washed, and incubated with 5 µM FCCP (Sigma-Aldrich, C2920) for 30 min to disrupt the membrane potential and finally observed at several time points to determine the differences in recovery between WT-HepG2 and Z-HepG2. To visualize lipid accumulation, HepG2 cells were incubated with Bodipy 493/503 (D3922; Invitrogen) at a final concentration of 2 µM. All experiments were performed in triplicate.

Immunofluorescence assays were performed using a STELLARIS Confocal Microscope (Leica Microsystems). After capturing the images, the staining was quantified using the CellProfiler software.

### Transmission Electron Microscopy (TEM)

For ultrastructural analysis, pellets of WT and Z HepG2 cells, as well as MM and ZZ organoids, were harvested by centrifugation and washed twice in Na2HPO4 (0.1 M) buffer (pH 7.4) buffer (phosphate buffer) at 4°C. Subsequently, they were chemically fixed with a mixture of 1% glutaraldehyde and 2% paraformaldehyde in phosphate buffer for 2 h at 4°C. Cells and organoids were further processed for electron microscopy following the same protocol. After fixation, they were washed three times in phosphate buffer at 4°C. Post-fixation was performed using osmium tetroxide (1%) and potassium ferricyanide (1%) in phosphate buffer for 1 h at 4°C and 0.15% tannic acid in phosphate buffer for 1 min at room temperature. samples were dehydrated in increasing concentrations of ethanol (50, 75, 90, 95, and 3 times with 100%) for 10 min each at 4°C. Infiltration was performed at room temperature with agitation using increasing concentrations of epoxy-resin (25, 50, 75, and 100%), and polymerization was performed at 60°C for 48h. Ultrathin sections of the samples (50-70 nm) were stained following standard procedures with 4% aqueous uranyl acetate and 2% lead citrate. The sections were then imaged using a FEI Tecnai-12 transmission electron microscope equipped with a LaB6 filament operated at 120 kV. Images were recorded at nominal magnification using a CCD (Charged Coupled Device) FEI Ceta digital camera.

### Seahorse assays

Differences in mitochondrial metabolism between the Z-AAT and M-AAT models were assessed by measuring oxygen consumption rates (OCR) and extracellular acidification rate (ECAR) using the Seahorse XF HS Mini Analyzer (Agilent). HepG2 cells and dissociated organoid cells were seeded on a seahorse plate at the required density (15.000-30.000 cells per well) and incubated at 37°C with 5% CO2 for 24 h. On the day of the assay, the medium was changed to seahorse medium (Seahorse XF DMEM 1X, 2 mM glutamine, 1 mM pyruvate, 10 mM glucose) and cell mitochondrial metabolism was analyzed using the Seahorse XF Cell Mito Stress Test Kit (Agilent) following the manufacturer’s recommendations. This type of test involves the injection of 1.5 µM oligomycin, 2.2 µM FCCP, and 0.6 µM rotenone/antimycin A (Rot/AA) into each well, which allows the OCR and ECAR to be measured under each condition, making it possible to calculate cells’ spare respiratory capacity, maximal respiratory capacity and ATP production. To allow data normalization, nuclei were stained with Hoechst (Invitrogen, #H3570) at a 1/300 dilution to count the number of cells per well. The results were analyzed using the Agilent Seahorse Analytics (Agilent) and GraphPad Prism software. In addition, mitochondrial metabolism was assessed by injecting three inhibitors of different pathways. To check whether mitochondrial metabolism was independent of the glucose pathway, the inhibitor UK5099 (Sigma-Aldrich, #5.04817.001) was injected at a final concentration of 2 µM. Similarly, to examine mitochondrial metabolism with the inhibition of the fatty acid oxidation pathway, etomoxir (Sigma-Aldrich, #236020) was injected at a final concentration of 4 µM. Finally, 3 µM BPTES (Sigma-Aldrich, #SML0601) was used to inhibit the glutamine pathway. Normalization and analysis were performed as previously described.

### Lipid supplementation

For lipid supplementation treatment, control and Z-HepG2 cells were seeded on a 6-well plate and treated with lipid concentrate (GibcoTM, #11905-031) at 1/300 dilution 24 hours prior to RNA extraction.

### Statistical analysis

Statistical analysis of the data was done using Excel and GraphPad Prism 10 Software to estimate the significance of the differences found between WT- and Z-HepG2, as well as MM and ZZ HepORG, in different assays. Results are presented as mean values ± SEM. Student’s t-test or Mann-Whitney U test was performed to determine statistical significance at p-values < 0.05.

### Study approval

All research was conducted in accordance with both the Declarations of Helsinki and Istanbul. Written informed consent was obtained from all participants, and the study was approved by the Ethics Committee of the Institute of Health Carlos III (Madrid, Spain).

## Data availability

Transcriptomics RNASeq data are deposited at ENA (European Nucleotide Archive) with data identifier PRJEB112082 for the patient-derived hepatic organoids, and PRJEB112083 for the Z-HepG2 and control cells. Proteomics data have been deposited at the PRIDE database with the dataset identifier XXXXXXX. All deposited data are publicly available as of the date of publication. Values for all data points in graphs are reported in the Supporting Data Values file. Other data will be shared upon reasonable request to the corresponding author.

## AUTHOR CONTRIBUTIONS

SGM, NM, OHD, EB, SM, JM, FD, SRS and MFP performed the experiments. SGM, NM, SPL and SJ wrote the draft of the paper. CBB, DM and MCT participated in result analysis. SGM, NM and SPL designed figures. AO, MR, MC, JRH gave clinical support and reviewed the manuscript. GGM, MJB, JA, and SJ contributed to the discussion and reviewed the manuscript. BMD coordinated the writing and critically reviewed and edited the manuscript.

## AKNOWLEDGEMENTS

We thank the BioNER -Biobanco Nacional de Enfermedades Raras- for their support in managing the samples.

